# Transmural and rate-dependent profiling of drug-induced arrhythmogenic risks through in silico simulations of multichannel pharmacology

**DOI:** 10.1101/752998

**Authors:** Ping’an Zhao, Pan Li

## Abstract

**Background:** In vitro hERG blockade assays alone provide insufficient information to accurately discriminate “safe” from “dangerous” drugs. Recent studies have suggested that the integration of multiple ion channel inhibition data can improve the prediction of drug-induced arrhythmogenic risks. In this study, using a family of cardiac cell models representing electrophysiological heterogeneities across the ventricular wall, we quantitatively evaluated transmural and rate-dependent properties of drug-induced arrhythmogenicity through computer simulations of multichannel pharmacology.

**Methods and Results:** Rate-dependent drug effects of multiple ion channel inhibition on cardiac electrophysiology at their effective free therapeutic plasma concentrations (EFTPCs) were investigated using a group of in silico cell models (Purkinje (P) cells, endocardial (Endo) cells, mid-myocardial (M) cells and epicardial (Epi) cells). We found that (1) M cells are much more sensitive than the other cell types to drug-induced arrhythmias and can develop early afterdepolarization (EAD) in response to bepridil, dofetilide, sotalol, terfenadine, cisapride or ranolazine. (2) Most drug-induced adverse effects, such as pronounced action potential prolongations or EADs, occur at slower pacing rates. (3) Although most drug-induced EADs occur in M cells, the application of quinidine at its EFTPC can cause EADs in all four cell types. (4) The underlying ionic mechanism of drug-induced EADs differs across different cell types; while I_NaL_ is the major depolarizing current during the generation of EAD in P cells, I_CaL_ is mostly predominant in other cell types. (5) Drug-induced AP alternans with larger beat-to-beat variations occur at high pacing rates in mostly P cells, while the application of bepridil can cause alternating EAD patterns at slower pacing rates in M cells.

**Conclusions:** In silico analysis of transmural and rate-dependent properties using multichannel inhibition data can be useful to accurately predict drug-induced arrhythmogenic risks and can also provide mechanistic insights into drug-induced adverse events related to cardiac arrhythmias.

**Author summary:** In vitro hERG blockade assays alone provide insufficient information to accurately discriminate “safe” from “dangerous” drugs, and computer simulation of ventricular action potential using multichannel inhibition data could be a useful tool to evaluate drug-induced arrhythmogenic risks. Our study suggested that the profiling of drug-induced transmural heterogeneities in cellular electrophysiology at all physiological pacing frequencies can be essential for the comprehensive evaluation of drug safety, and for the quantitative investigation into ionic mechanisms underlying drug-specific arrhythmogenic events. These in silico models and approaches may contribute to the ongoing construction of a comprehensive paradigm for the evaluation of drug-induced arrhythmogenic risks, potentially increase the success rate and accelerate the process of novel drug development.

## Introduction

Drug-induced cardiotoxicity has been a major concern since the early stage of novel drug development. Unexpected post-marketing occurrence of cardiotoxic effects remains a leading cause of drug withdrawal and relabelling^[1–3]^. As defined by the International Conference of Harmonization Expert Working Group for all drugs in development, QT interval prolongation has been used as a biomarker to predict the potential risk of Torsade de Pointes (TdP)^[4, 5]^. Most drugs that prolong the QT interval inhibit cardiac potassium channels encoded by human ether-à-go-go related gene (hERG); therefore, the level of hERG channel inhibition has been the “gold standard” to predict the TdP risk. However, recent studies suggested that the in vitro hERG blockade assay alone provides insufficient information to accurately discriminate “safe” from “dangerous” drugs. For instance, QT prolongation can be induced by drugs that inhibit other ionic channels such as I_Ks_, and it has been known for years that the arrhythmia associated with hERG blockade is mitigated by concurrent blockade of Na^+^ or Ca^2+^ channels^[6, 7]^. Other electrocardiogram (ECG) abnormalities, such as QT shortening or T wave alternation, are also frequently associated with TdP. Recently, the Comprehensive in vitro Proarrhythmia Assay (CiPA) has been proposed to address the misidentification issue of drug-associated TdP risk based on hERG inhibition and QT prolongation data. This new paradigm is based on integrated assessment of multiple ion channel dynamics in delayed ventricular repolarization; alterations to this process lead to repolarization in stability and arrhythmias^[8]^.

Computational modelling of the heart has been an important tool in advancing our understanding of cardiac excitation-contraction coupling. Recent studies have utilized mathematical modelling to evaluate the drug-induced proarrhythmic risk^[9–12]^. For example, multiple cardiac ion channels were integrated into the human myocyte model to improve the assessment of proarrhythmic risk^[13]^. Additionally, research institutions classify the cardiac toxicity of drugs through a computational approach that combines a human ventricular myocyte model of drug effects and machine learning^[14]^. However, researchers have performed experiments on only one cell type. Here, we employed a series of mathematical models and performed quantitative analyses of drug-induced arrhythmogenic risk in four cell types and physiological frequencies through multichannel pharmacology.

## Results

### Transmural heterogeneity of cardiac AP morphologies and adaptations

Transmural action potential (AP) morphologies are shown during steady-state pacing (cycle length (CL)=1000 ms) and action potential duration (APD) rate adaptations (Figure 1). The APD in mid-myocardial (M) cells was longer than that in epicardial (Epi) cells or endocardial (Endo) cells and was considerably shorter than the APD in Purkinje (P) cells. The AP amplitude in P cells was considerably higher than that in different types of ventricular cells. In Epi and M cells, AP reproduced a characteristic phase 1 notch and dome morphology, which was not apparent in Endo cells. In addition, the notch and dome morphology was absent in simulated P cells. The APD was prolonged as the heart rate slowed down. The simulated cell AP morphology and APD adaptation curve were consistent with previous experiments^[18, 19]^.

**Fig 1.**
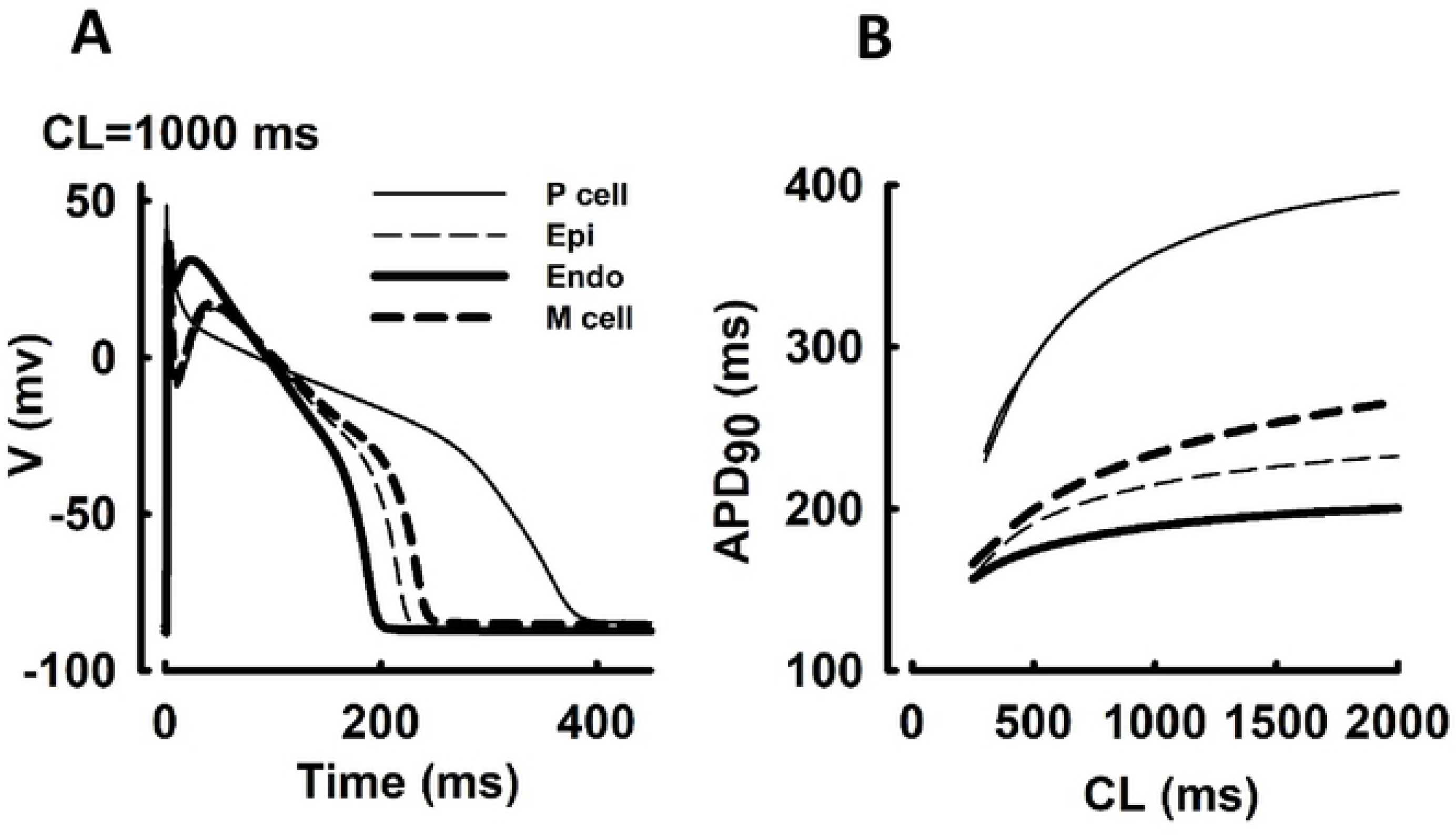
Transmural AP morphologies (A) and APD rate adaptations (B) in Purkinje (P), epicardial (Epi), endocardial (Endo) and mid-myocardial (M) cells.

### Drug-induced changes in AP adaptations

Drug-induced changes in AP adaptations were shown in Figure 2. For bepridil, cisapride, terfenadine and dofetilide, simulated APD was significantly prolonged (the percentage of prolongation>10%) in all cell types and moderately prolonged for sotalol and ranolazine, and EAD was triggered exclusively in M cells. In contrast, chlorpromazine, verapamil and ondansetron caused moderate prolongation, and diltiazem and mexiletine led to minor APD shortening by 0.9% and 2.6% at CL=2000 ms in P cells, respectively, while causing a positive shift of APD adaptation curves in other cells. Furthermore, at CL=1000 ms, cisapride caused APD prolongation of 12.6%, 14.3% and 15.7% in P, Epi and Endo cells, respectively, while it caused APD prolongation of 129.3% in M cells. In general, the percentages of drug-induced APD prolongations were much higher in M cells. For example, at CL=1000 ms, bepridil caused APD prolongation of 15% (P cells), 27% (Epi cells) and 25.6% (Endo cells) respectively, and caused APD prolongation of 158% in M cells. Drug-induced changes in APD is highly dependent on pacing CLs or rates, with drug-induced EAD events mostly associated with slower pacing rates. For example, in M cells, the percentage of APD prolongation of ranolazine was 7.5% (CL=500 ms) and 9.4% (CL=1000 ms), while ranolazine induced EAD (102.9%) at CL=2000 ms. These results suggested that M cells could be more sensitive in general to drug-induced APD prolongation and EAD events, and interestingly P cells are more sensitive to AP alternans.

**Fig 2.**
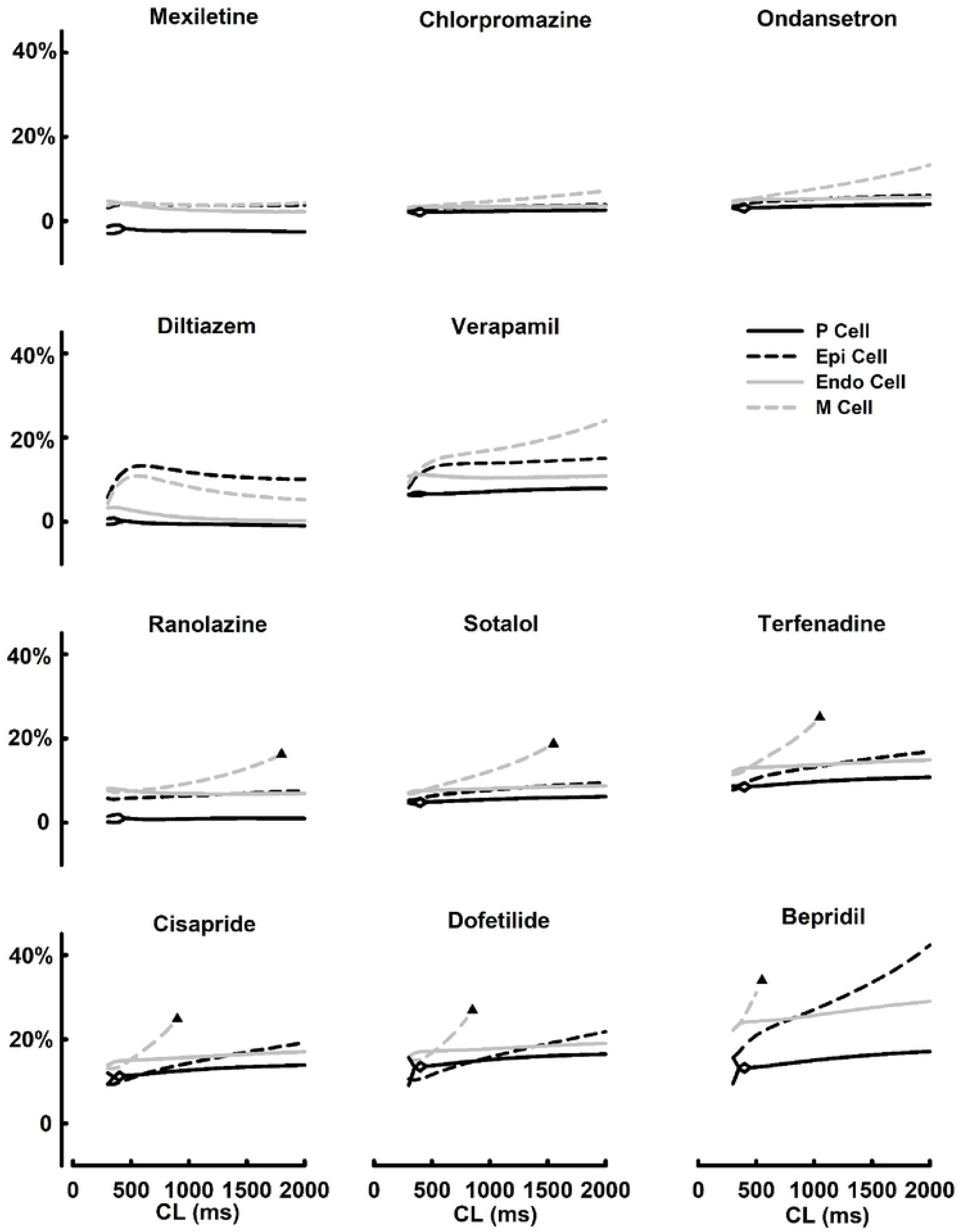
Drug-induccd changes in AP adaptations in Purkinje (P), endocardial (Endo), mid-myocardial (M), and epicardial (Epi) cells. All drugs were applied at their effective free therapeutic plasma concentrations (EFTPCs), and black triangles indicate occurances of EAD events.

### Transmural characteristics of drug-induced arrhythmogenicity

The AP and ionic channel dynamics of cisapride and dofetilide were examined at fixed BCL =1000 ms (Figure 3). In Endo cells, cisapride caused APD prolongation of 15.7%, and the inward current of I_NaL_ displayed a slight increase compared to that of the control. The inward current of I_CaL_ and the outward current of I_to_ were almost the same as those of the control. The outward current of I_Kr_ displayed a 29.3% reduction in the peak. In M cells, drug-induced changes in I_NaL_ and I_Kr_ were mostly responsible for APD prolongation of 129.3%. I_to_ was larger in M cells than in Endo cells, resulting in a significantly greater notch V_max_ in M cells. I_CaL_ plays a major role in the genesis of EADs, and the reactivation of I_CaL_ coincided with EADs (Figure 3A). In P cells, dofetilide caused APD prolongation of 15.1%, while it was 132.6% with EAD generation in M cells. In M cells, EAD occurs because I_CaL_, I_NaL_ and I_Kr_ play significant roles in the prolongation of APD (Figure 3B).

**Fig 3.**
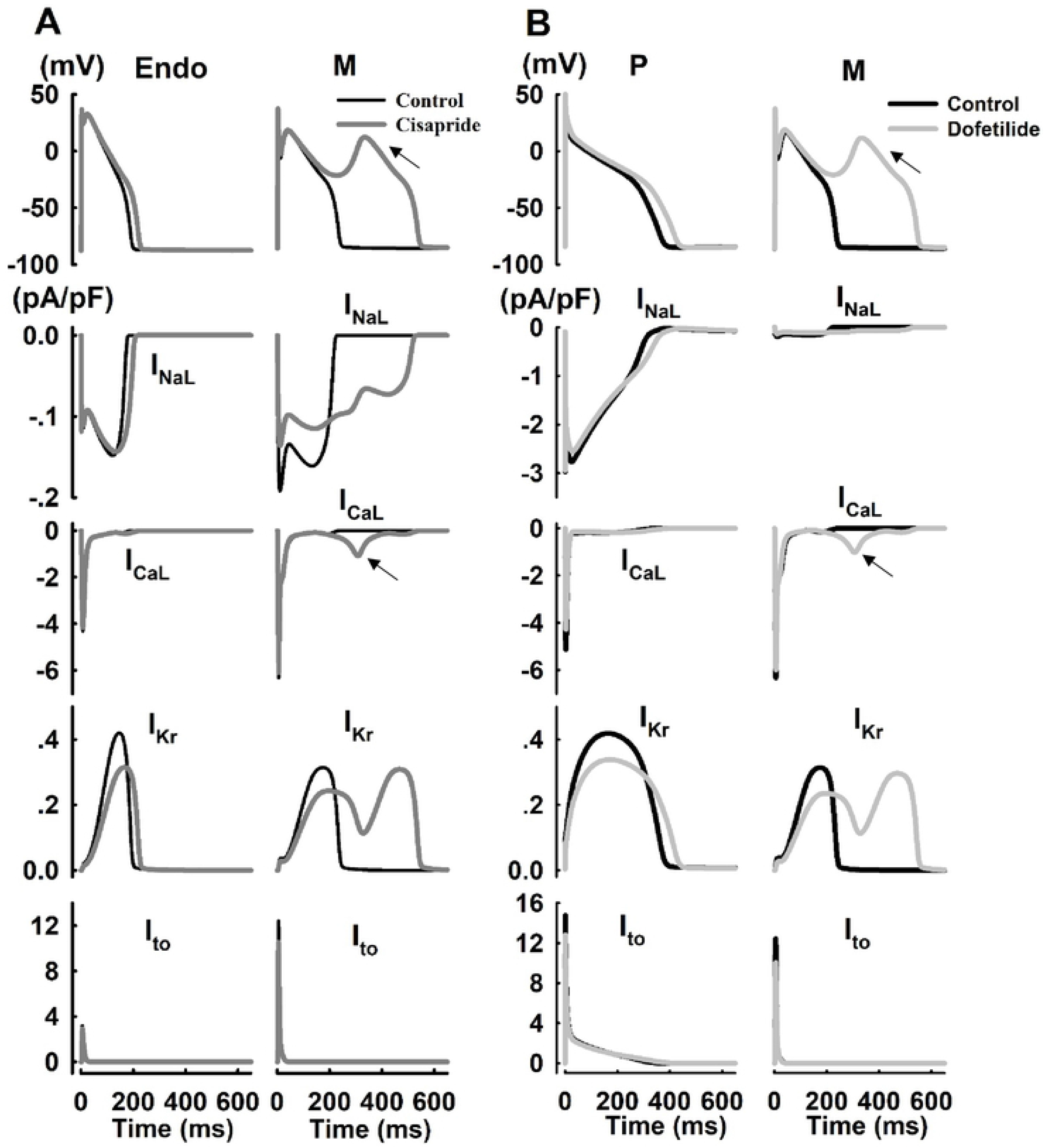
Cell dependent properties of cisapride and dofetilide at fixed CL=1000 ms. A: APs and ionic currents (I_NaL_, I_CaL_, I_kr_, and I_to1_) in Endo and M cells with the application of cisapride; B: APs and ionic currents (I_NaL_, I_CaL_, I_kr_, and I_to1_) in P and M cells with the application of dofetilide.

### Rate-dependent properties of drug-induced arrhythmogenicity

To assess the rate dependence, we evaluated ionic channel dynamics at different frequencies (0.5Hz,1Hz, and 2Hz) with the application of ranolazine and bepridil in M cells. Ranolazine caused an APD prolongation of 7.5% (CL=500 ms), 9.4% (CL=1000 ms) and 102.9% (CL=2000 ms), respectively. At CL=500 ms, hERG blockade was mitigated by the concurrent inhibition of Na^+^ channels (I_NaL_), resulting in slight prolongation of APD. At CL=2000 ms, in addition to the interplay between I_Kr_ and I_NaL_, the inward current I_CaL_ increased, giving rise to the pronounced prolongation of APD and occurrence of EAD (Figure 4A). Bepridil caused APD prolongation of 31%, 158.1% and 255.2%, at CL=500 ms, 1000 ms, and 2000 ms, respectively. At CL=500 ms, I_Kr_ was a major current responsible for APD prolongation; inhibition of I_NaL_ could shorten APD, attenuate the prolongation resulting from hERG (I_Kr_) inhibition and prevent EAD (Figure 4B).

**Fig 4.**
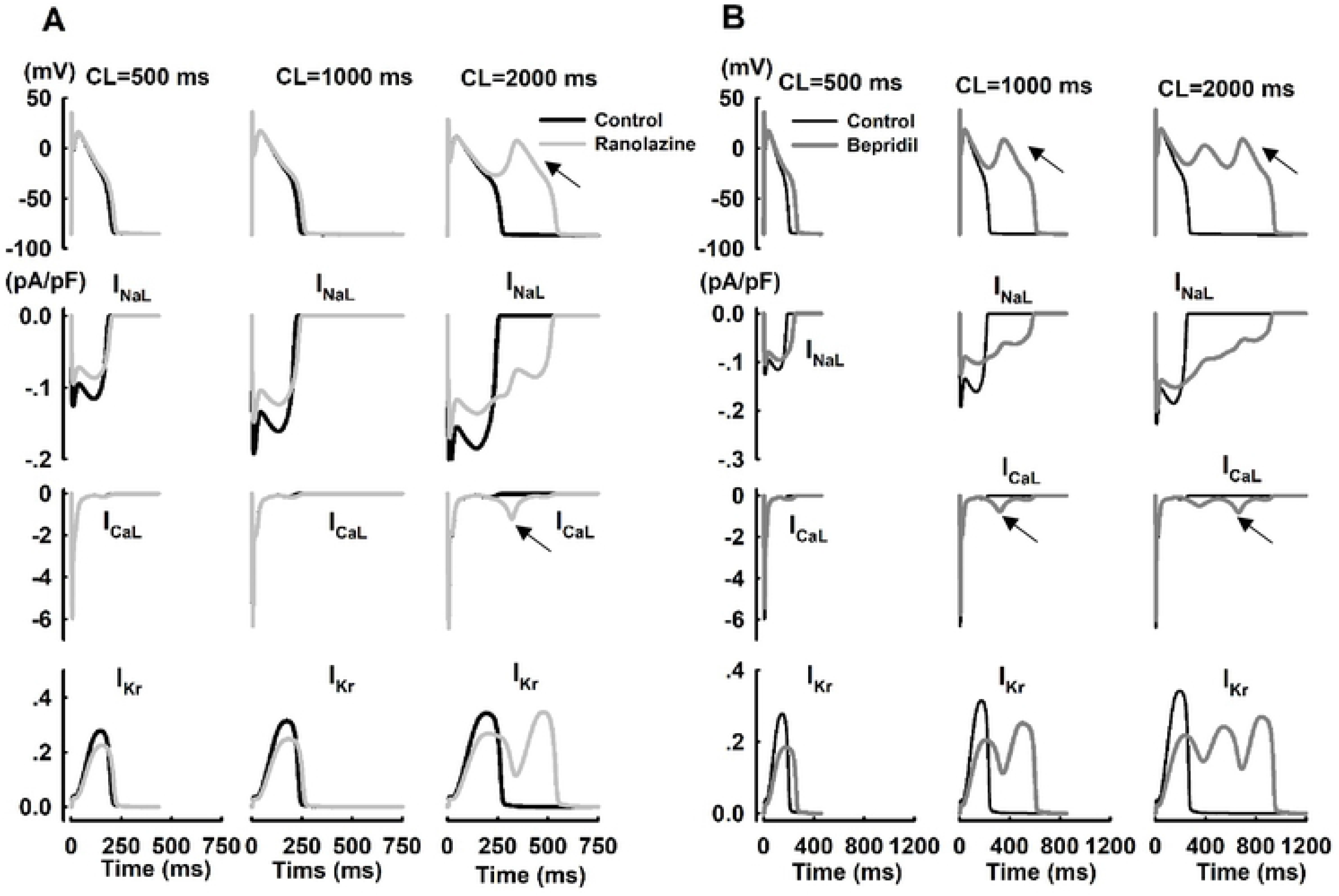
Rate-dependent properties of ranolazine and bepridil in M cells at different pacing CLs. A: APs and ionic currents (I_NaL_, I_CaL_, and I_kr_) at different CLs with the application of ranolazine; B: APs and ionic currents (I_NaL_, I_CaL_, and I_kr_) at different CLs with the application of bepridil.

### Drug-induced early afterdepolarization (EAD) and AP alternans

With the application of quinidine in all four types of cells at CL=2000 ms, large amplitude EADs developed starting from the 133^rd^ beat in Endo cells (Figure 5A). Persistent EAD events occurred starting at the 4^th^ beat in M cells, and these cells died at the 6^th^ beat (Figure 5B). EADs developed starting from the 3rd beat in Epi cells (Figure 5C). Consecutive EADs were generated from the 1503^rd^ beat (Figure 5D). Additionally, during the generation of EAD, I_CaL_ produced a current of much larger amplitude than did I_NaL_, and I_CaL_ preceded the reactivation of I_NaL_ in ventricular cells (Endo, M, and Epi). However, in P cells, the reactivation of I_NaL_ was larger in amplitude and preceded that of I_CaL_. I_CaL_ also reactivated during the EAD upstroke, but its contribution was secondary to that of I_NaL_. Thus, the ionic mechanisms of EAD in P cells or V cells rely on different ion channels during a prolonged plateau; I_NaL_ is the major depolarizing current for EAD in P cells, while in V cells, it is I_CaL_.

**Fig 5.**
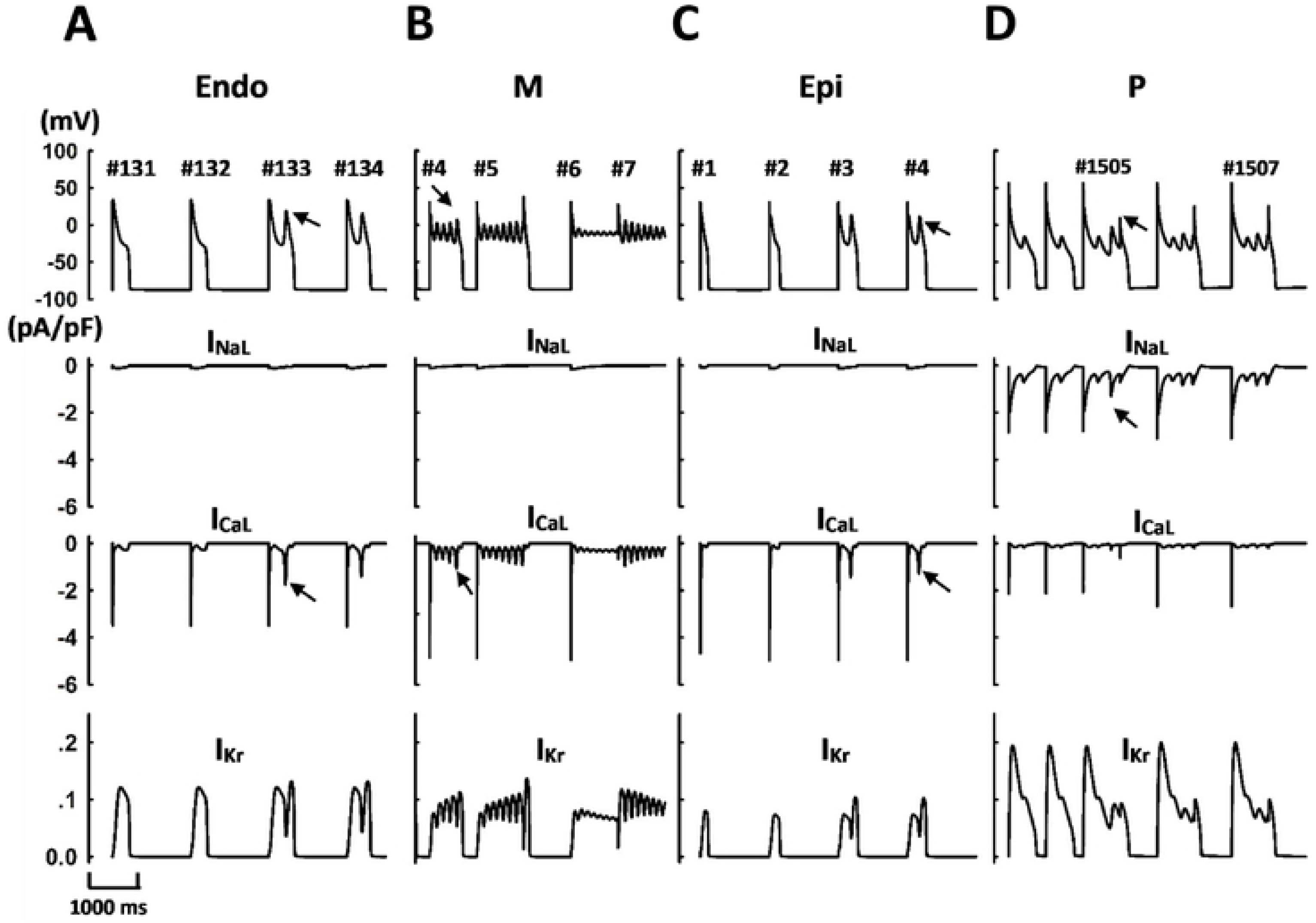
EAD events and underlying ionic currents with the application of quinidine in Endo (A), M (B), Epi (C) and P (D) cells at a fixed CL=2000 ms.

Drug-induced AP alternans and underlying ionic currents were shown in Figure 6. With the application of dofetilide or bepridil, AP prolongation with enhanced beat-to-beat variations (alternans) was observed at CL=300 ms in P cells; with the application of verapamil, AP prolongation with no beat-to-beat variations was shown. In addition, the application of bepridil can also cause alternating EAD patterns at slower pacing rates (CL=1050ms) in M cells. While major reduction in I_Kr_ was most responsible for drug-induced APD prolongation, a smaller I_to_ could play the secondary role with the application of dofetilide (Figure 6B). Furthermore, it was shown that I_NaL_ and I_CaL_ could be the major currents responsible to induce AP alternans, and I_ks_ and I_NCX_ may function as protecting currents against AP alternans (Figure 6B and 6C).

**Fig 6.**
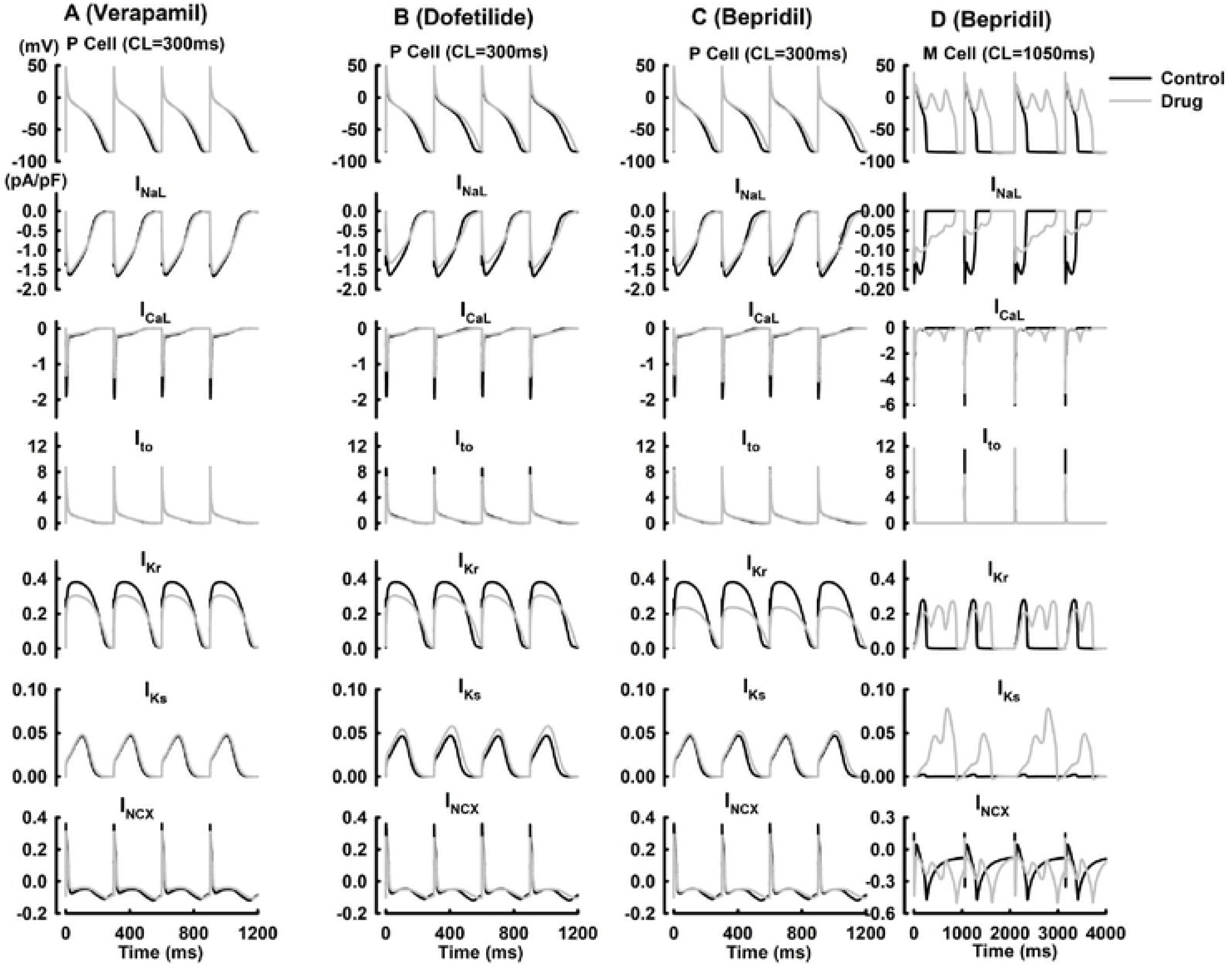
AP alternans and underlying ionic currents with the application of verapamil (A), dofetilide (B), bepridil (C) in P cells at CL=300ms, and bepridil in M cells at CL= 1050ms (D).

## Discussion

In this study, we quantitatively evaluated the transmural characteristics and rate dependence of drug-induced arrhythmogenicity through simulations of multichannel pharmacology using a family of cardiac cell models. To address the misidentification of drug-associated TdP risk based solely on hERG and QT data, Kramer et al.^[20]^ published a predictive model that accounts for L-type Ca^2+^channel blockade in addition to hERG blockade. This model improved discrimination between torsadogenic and nontorsadogenic drugs over the hERG assay, and a new model that combines dynamic drug-hERG interactions and multichannel pharmacology was developed that improved early prediction of compounds’ clinical torsadogenic risk^[13]^. In our study, the drug-induced arrhythmogenic risk was evaluated in diverse cell types. We found that M cells, in general, are much more vulnerable to drug-induced AP prolongation and EAD generation, which suggest that the intrinsic arrhythmogenicity of M cells can be much higher than that of other cell types. In addition, we found that drug-induced changes in APD adaptation can be important during the evaluation of drug-induced cardiac toxicity. The pronounced increase in APD at slower heart rates allows for the recovery of inactivated calcium or sodium channels, widening the window of EAD generation, a cellular event that can induce TdP^[21–24]^. Furthermore, we found that quinidine can generate EAD in all cell types, and the predicted high TdP risk of quinidine is consistent with that in previous experimental and clinical studies. However, the mechanism of drug-induced EAD generation differs across cell types; EAD generation in P cells is mostly due to reactivation of I_NaL_, while in Endo, M and Epi cells, I_CaL_ plays the predominant role^[15]^. In addition, we found that comparing to other cell types, P cells are generally more sensitive to drug-induced changes in AP alternans at fast pacing rates, that can potentially lead to Purkinje-ventricular conduction abnormalities at tissue/organ level. Interestingly, with the application of bepridil, AP alternates with beat-to-beat occurrences of EAD events can be observed in M cells at slower pacing rates. These in silico findings support our hypothesis that it may be insufficient to evaluate drug-induced arrhythmogenic risk in any single cell type and at a given frequency.

We concluded that simulations of multichannel pharmacology in diverse cell types at all physiological pacing rates are essential to evaluate drug-induced arrhythmogenic risks. However, the heart is a complex biological system, and our study was limited at the cellular level, with no evaluation of tissue or organ level complexities. For example, a recent study presented an arrhythmic hazard map for a 3D whole-ventricle model under multiple ion channel inhibition to predict drug-induced arrhythmogenic risks. However, one advantage of the series of models used in our study is its computational efficiency, which may enable large-scale in silico screening for compounds of high cardiac safety, and these quantitative approaches could also offer further mechanistic insights into drug-induced arrhythmogenic risk.

## Methods

P, Endo, M, and Epi cell models were derived from the Pan-Rudy(PRd)^[15]^ and Keith-Rudy (KRd)^[16]^ models based on experimental data on transmural heterogeneity in electrophysiology by altering G_NaL_, G_to1_, G_Ks_ and G_naca_ (see Supplemental Material Table 1 for details). We simulated the application of 12 drugs ^[17]^ at their EFTPCs on seven ion channels (I_Na_, I_NaL_, I_CaL_, I_Kr_, I_Ks_, I_to_ and I_K1_) in four cell models (see Supplemental Material Table 2 for details). APD was defined as the duration from maximum dVm/dt during the AP upstroke to 90% of full repolarization (APD_90_). All cell simulations were paced for 60 min to reach steady states.

To simulate a drug blocking a channel, we used the following equation to scale gx, based on the drug IC_50_ and concentration:

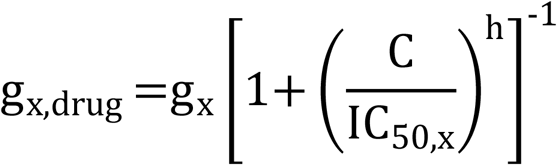

where gx, drug is the maximal conductance of channel x in the presence of the drug; C is the concentration of the drug; IC_50,x_ is the half-maximal inhibitory concentration for that drug and current through channel x; and h is the hs of the drug.

## Supporting information

**S1 Table 1.** Model details of Epi, Endo and M cells according to experimental measurements.

**S1 Table 2.** IC50s and Hill coefficients (h) of drugs were calculated using the inhibition data from the study by Crumb et al.

## Acknowledgments

This work was supported by the National Science Foundation of China (Grant No. U1604178 to P.L.).

